# The circulatory physiopathology of human red blood cells investigated with a multiplatform model of cellular homeostasis. II. Capillary transits and the role of PIEZO1

**DOI:** 10.1101/2020.03.07.981787

**Authors:** Simon Rogers, Virgilio L. Lew

## Abstract

In this and the next paper of this series we apply the red cell model introduced in the previous paper to investigate the changes in RBC homeostasis during capillary transits and over the full circulatory lifespan of the cells. These are topics inaccessible to direct experimentation but rendered mature for a modelling approach by recent findings and by a large body of apparently unrelated early results which robustly constrain the parameter space offering the opportunity for an in depth study of the mechanisms involved. Capillary transit times vary between 0.5 and 1.5s during which the red blood cells squeeze and deform in the capillary stream transiently opening stress-gated PIEZO1 channels, creating minuscule quantal changes in RBC ion contents and volume. Widely accepted early views originally based on results from experimentally shear-stressed red cells suggested that quantal changes generated during capillary transits add up over time to generate the documented changes in RBC density during their long circulatory lifespan, the quantal hypothesis. Applying the new PIEZO1 extension of the RBC model (RCM) introduced in the previous paper we investigated in detail the changes in homeostatic variables that may be expected during single capillary transits resulting from transient PIEZO1 channel activation. The predicted quantal volume changes were infinitesimal in magnitude, biphasic in nature, and essentially irreversible within inter-transit periods. A sub-second transient PIEZO1 activation triggered a sharp swelling peak followed by a much slower recovery period towards lower-than-baseline volumes. The peak response was caused by net CaCl_2_ and fluid gain via PIEZO1 channels driven by the steep electrochemical inward Ca^2+^ gradient. The ensuing dehydration followed a complex time-course with sequential, but partially overlapping contributions by KCl loss via Ca^2+^-activated Gardos channels, PMCA mediated calcium extrusion and chloride efflux by the Jacobs-Steward mechanism. The change in relative cell volume predicted for single capillary transits was below 10^−4^, an infinitesimal volume change incompatible with a functional role in capillary flow. The biphasic response predicted by the RCM appears to conform to the quantal hypothesis, but whether its cumulative effects could account for the documented changes in density during RBC senescence required an investigation of the effects of myriad transits over the full period of circulatory lifespan, the subject of the next paper of this series.

## Introduction

The RBC-mediated transport of O_2_ and CO_2_ between lungs and tissues evolved to operate with minimal energy cost to the organism. A first critical feature enabling such economy is the extremely low cation permeability of the RBC membrane [1–3]. This allows the cells to maintain steady volumes for extended periods of time with minimal cation traffic, pump-leak turnover rates and ATP consumption. Glycolytic ATP turnover by the full RBC mass of a healthy human adult amounts to less than 0.06% of the total body ATP turnover [4].

A second critical feature of the optimized economy concerns the compromise between RBC turnover rate and circulatory lifespan. RBCs are the most abundant cells in the body, their mass adapted for adequate gas transport at all levels of physiological demand. Biosynthetic and biodegradable replacement of such a large cell mass imposes a heavy metabolic cost to the organism which can only be reduced by extending the circulatory lifespan of the cells thereby reducing their replacement frequency. Circulatory longevity, on the other hand, is limited by the extent to which RBCs, without nucleus and organelles, and devoid of biosynthetic capacity, can sustain the functional competence of its metabolic and membrane transport components required for volume stability and optimal rheology. Optimal rheology requires that the RBCs retain a large degree of deformability for rapid passage through narrow capillaries and for ensuring minimal diffusional distances for gas-exchange across capillary walls. Deformability, in turn, depends on the RBCs maintaining their volume well below their maximal spherical volume (reduced volume), a condition fulfilled when their reduced volume is kept within a narrow margin around 60% [5]. For human RBCs with a mean circulatory lifespan of about 120 days, this represents a substantial challenge, and the mechanism by which this is achieved is the subject of the investigation in this and the next paper [6].

RBCs undergo gradual densification for most of their lifespan [7, 8]. After about 70-90 days, the densification trend reverses [9–12], a phenomenon interpreted as a strategy evolved to prolong the circulatory survival of the cells by keeping them within the optimal reduced volume range, otherwise imperilled by sustained dehydration. The terminal clock is activated by immune signalling at the RBC surface via the Kay-Lutz mechanism eliciting RBC phagocytosis by spleen and liver macrophages [13–16]. There is also solid evidence documenting age-dependent late reductions in the activity of the Na/K pump (ATP1A1 gene) [17–19], gradual decline in Ca^2+^ pump activity (PMCA, PMCA4b, ATP2B4 gene) [20–23], and in Gardos channel activity (KCNN4) [24] among membrane transporters, and in the activities of other enzymes and membrane components [7], reductions generally attributed to unavoidable damage and decay in a cell deprived of protein renewal capacity.

The involvement of increased calcium permeability and Gardos channels in RBC dehydration has been documented in many haematological pathologies and in a wide range of experimental conditions. Their participation has also been frequently suggested to set the pattern of the homeostatic changes during circulatory aging despite lack of reliable early evidence of increased Ca^2+^ permeability states in physiological conditions. Following pioneering work in the eighties suggesting that shear stress triggered the progressive dehydration of RBCs in the circulation [25, 26], major progress in this area was made with the discovery by Dyrda et al., [27] that local membrane deformations activate Ca^2+^-dependent K+ and anionic currents in intact human RBCs, and subsequently by the finding that RBC membranes express the stress-activated PIEZO1 channel [28–33], a prime candidate pathway to account for the observed deformation-induced increase in Ca^2+^ permeability [27, 34]. More recently, elegant experiments with Fluo-4-loaded RBCs passing through constrictions in microfluidic chips have provided firm evidence on the deformation-calcium signalling link, its reversibility and time-course [35].

These findings have stimulated intense research on the participation of PIEZO1, Gardos channels, and of mutant forms of these transporters in normal RBC homeostasis and in inherited pathologies of RBC hydration. Because PIEZO1 channels are poorly ion selective [36], each capillary passage would be expected to trigger gradient dissipation for K^+^, Na^+^, Ca^2+^, Mg^2+^ and Cl^−^, creating minute changes in RBC ion contents. Because of its huge inward electrochemical gradient even a minor and brief increase in Ca^2+^ permeability could generate a significant elevation in [Ca^2+^]_i_ above baseline levels, a signal with the power to activate Ca^2+^-sensitive Gardos channels with downstream effects of KCl loss and cell dehydration. The cumulative effect of such quantal steps during capillary transits was assumed to drive the gradual densification of RBCs during most of their circulatory lifespan, the quantal hypothesis [4, 25].

We investigate here the changes in RBC variables predicted for single capillary transits applying a newly introduced PIEZO routine integrated in the JAVA version of the RBC model (RCM) [37]. The cumulative effect of repetitive capillary transits over the full circulatory lifespan of the cells was investigated in the accompanying paper [6]. At each stage in the course of this investigation compliance with reliable experimental data was used as the constraining guide on the range of acceptable model outcomes and parameter values.

## Methods

### Modelling strategy

From the start, it was critical to select the body of knowledge we wished the model outcomes to comply with, and to establish a hierarchical scale of weight for the different parameters based on experimental support and relevance. The experimental data chosen to constrain and guide this investigation was primarily based on relatively recent results by Dyrda et al., [27], Glogowska et al., [38], Danielczok et al., [35], Kuchel and Shishmarev, [34], Zhao et al., [36] Ganansambandam et al., [39], and also on dated results that harmonize with the more recent findings [40–42]. Five PIEZO1-attributed parameters defined the model protocol for single capillary transit simulations: duration of the open-state (OS, in s), three cationic permeabilities for Ca^2+^, Na^+^ and K^+^, PzCa, PzNa, PzK, respectively (in h^−1^), and one anionic permeability for the combined diffusible Cl^−^ + HCO_3_^−^ anions, PzA (h^−1^). It is important to stress here that the attribution of an anion conductance to PIEZO1 (PzA) is only intended as a means to implement a minimalist phenomenology that enables us to investigate the need or otherwise to associate the changes in PIEZO1-mediated cation and anion permeabilities during the PIEZO1 open state in the simulations. As will become apparent in this and the following paper [6], the results do support a strong association between increased cation and anion permeabilities during the PIEZO1 open states. However, the model results carry no information on whether the anion permeability is through PIEZO1, as PzA may imply, or through another PxA pathway. The association may be hypothetically explained by interactions between different channels. For instance, the kinetics of PIEZO1 and that of the slippage conductance through the anion exchanger, AE1 [43, 44] may be somehow linked through their connections to the underlying cytoskeletal mesh, generating a transient AE1 configuration with enhanced conductance during the PIEZO1 open state. Similar arguments may apply to other anionic conductance pathways described in the literature [27, 38, 45].There is no information on the actual value or interdependence of any of the PIEZO1 attributed parameters in the context of capillary transits, leaving this model exploration widely open.

Images of RBCs during capillary transits show RBCs squeezed, elongated and deformed in the flow, often in single filing trains. During these brief transits, of between 0.5 and 2s duration, the volume of plasma surrounding the cells becomes much smaller than the actual volume of the cells, a condition that can be represented in the model with cell volume fractions (CVF) near one, during which any PIEZO1-mediated change in cell contents will substantially affect the composition of the surrounding medium. During inter-transit periods, on the other hand, before RBCs ingress to, or after they emerge from capillaries, RBCs flow in the systemic circulation which acts like an infinite reservoir of a composition regulated by RBC-independent processes. Modelling such transit-inter-transit transitions requires a protocol which ensures that the composition changes within the miniscule volumes of medium surrounding the cells during the brief capillary transits are not carried over and accumulated in the medium surrounding the cells during the inter-transit periods in the systemic circulation. This required introducing a “restore medium” subroutine for inter-transit periods to minimize carryover effects. Because transit times are so short (< 2s) relative to inter-transit periods (1-2 min) it was expected that running a control at low CVF throughout would provide an adequate approximation to the results of realistic simulations, a necessary test to rule out unsuspected effects.

We carried out a large number of preliminary tests with the RCM, seeking compliance with documented experimental results. Four transport systems were found to be the main determinants of the response to deformation-induced changes in RBC homeostasis: PIEZO1 with PzA, the trigger of all downstream effects, Gardos channels, the PMCA and the Jacobs-Stewart mechanism (JS) ([37], Fig 7). The pattern that emerged from these trials and that fulfilled the expected compliance could be implemented with a rather restricted set of parameter values. To report in the simplest possible terms the process followed to arrive at this result, and to explain the mechanisms involved, we generated a standardized Reference protocol (Fig 1) which delivers the curves labelled “Ref” in the figures (in black). The effects of parameter variations are shown by comparison to Ref. The Ref protocol runs a standard baseline period of two min. Ingress of a RBC in a capillary is simulated with an instant CVF transition from 0.00001 to 0.9 together with PIEZO1 activation. After 0.4s, egress is simulated with instant PIEZO1 closure and with the RBC returning to CVF = 10^−5^ in a restored medium condition. The standard duration of post-transit periods designed here to provide adequate information on volume changes and trends was set to five minutes, not to be confused with inter-transit durations. The default permeability values set for the Ref routine were: PzCa, 70h^−1^; PzA, 50h^−1^; and zero for PzNa and PzK. The reasons for these choices will become clear with the analysis of the results in this paper; a full explanation for the value of 70h^−1^ attributed to the reference PzCa will have to await the analysis in the section on Hyperdense collapse in the next paper [6].

**Figure 1.**
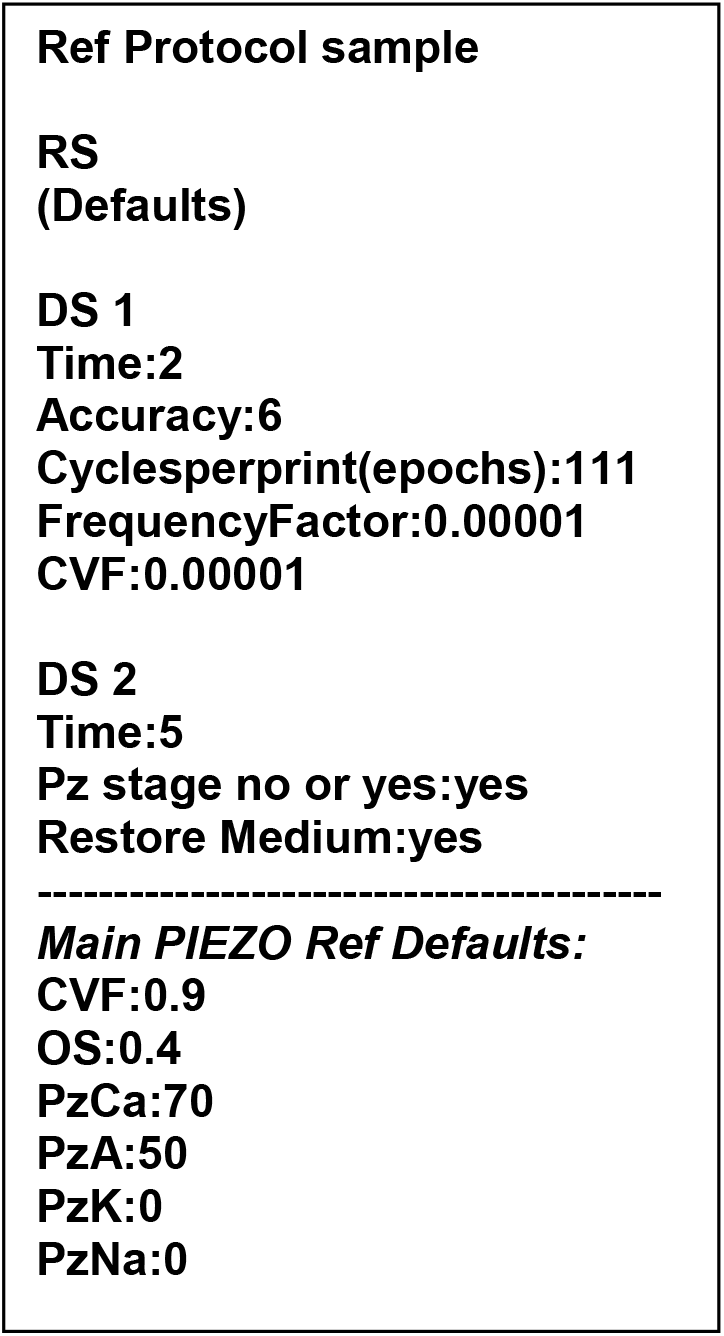
Default Reference protocol (Ref) for testing the effects of parameter and variable changes on RBC responses during and after capillary transits. The default protocol runs a three-step dynamic sequence on a RBC configured by the defaults in the Reference State (RS): 1) a two minute baseline, 2) at t=2min, ingress to a capillary is defined by an instant transition from a very low (CVF=10^−5^) to a very high (CVF=0.9) cell volume fraction together with opening of PIEZO1 channels for 0.4s, and 3) at t=2min+0.4s, an instant capillary egress transition back to a very low CVF (10^−5^) with PIEZO1 inactivation and medium composition restored to baseline values. The default duration of the post-transit stage within the PIEZO routine was set to 5min. Within the DS1 baseline period, we set the data output conditions to be followed through DS1 and DS2: “accuracy” was set to 6, the minimal value found to be required for reporting the miniscule size of the predicted changes; “Epochs” was set to 111 and “Frequency factor” to 10^−5^, values found to offer acceptable point densities in the model outcomes for discerning predicted time-dependent trends unambiguously. For the DS2 period covering the brief capillary transit and subsequent relaxation period back in the systemic circulation, we start by setting the overall duration of this stage (5min in the Ref sample) and then bring up the dedicated PIEZO routine confirming with “yes” the prompts on “Pz stage no or yes” and on “Restore Medium”. All defaults are open to change as extensively implemented in figures 2 and 4.

**Figure 2.**
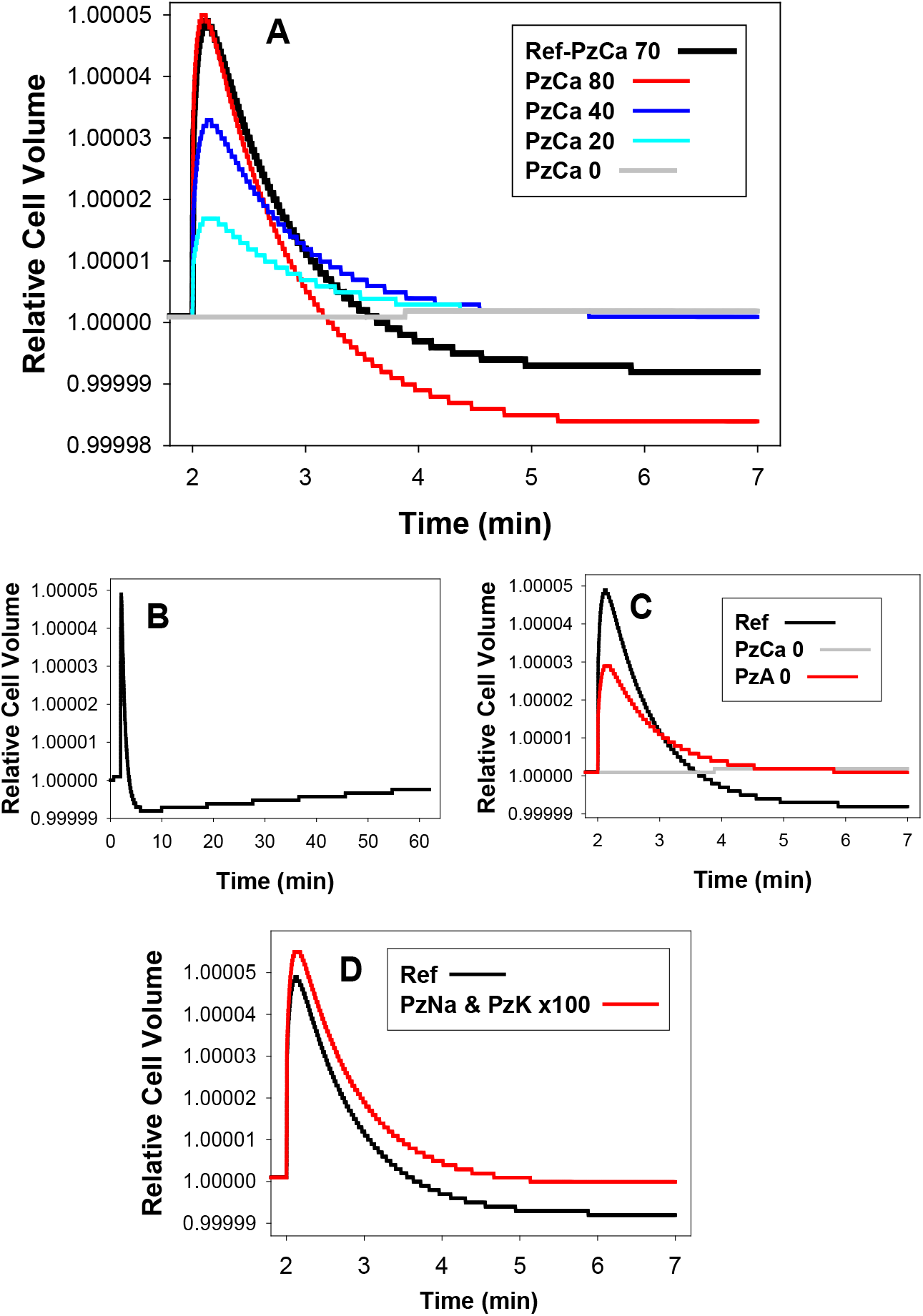
Predicted volume effects of PIEZO1 activation during capillary transits. After a baseline period of 2 min, ingress into capillaries at t = 2 min was simulated with a transition from a cell/medium volume ratio (CVF) of 0.00001 to 0.9 and a sudden opening of PIEZO1 channels activating electrodiffusional permeabilities to Ca^2+^ (PzCa), monovalent anions (PzA; A^−^ lumps Cl^−^ and HCO_3_^−^), and monovalent cations (PzNa and PzK), as indicated. Capillary egress was set simultaneous with PIEZO1 channel closure (all PzX = 0) 0.4 seconds after ingress, together with a return to a CVF of 0.00001. RCV changes were followed thereafter for different lengths of time. Reference curves (Ref) are shown in black in all panels and were simulated with defaults PzCa = 70h^−1^ and PzA=50h^−1^. **A**. Effects of different PzCa values. **B**. Reference condition followed for 60min to estimate longer-term reversibility of PIEZO1-elicited volume changes. **C**. Short- and long-term effects of PzA = 0 relative to Ref with PzA = 50/h; PzA differences in the range 30 to 50/h, the range of values found to fit observed activated anion conductances under on-cell patch clamp [27], had no significant effects on Ref. **D**. Effects of increasing PzNa and PzK permeabilities to values 100-fold above those of the basal RBC Na^+^ and K^+^ permeabilities.

Points noted during preliminary tests, of secondary relevance but considered necessary as background information, is succinctly summed up next. Interested readers are encouraged to design protocols and replicate these tests with the RCM. 1) It made little difference to the results whether RBCs were simulated emerging from the capillary immediately after PIEZO1 closure or after lingering in the high CVF conditions for up to 2s after PIEZO1 closure. This validated the use of simpler protocols with the same duration set for PIEZO1 open states and transit times (Fig 1). 2) PIEZO1 open state duration proved to be inversely related to permeabilities. For instance, the calcium influx generated by a PzCa of 70h^−1^ over 0.4s rendered similar results with PzCa of 140h^−1^ over 0.2s. Thus, OS duration and permeabilities are inversely related, not independent parameters in the context of their modelled effects. 3) Comparison of results between simulations with CVF values of 0.9 and 0.00001 during transits showed substantial agreement, as anticipated because of the brevity of the capillary transit periods (not shown). Nevertheless, the choice was made to retain the more realistic protocols with high-low CVF transitions in the presentation of the results, as differences of negligible size at a single transit level were shown to have significant effects when accumulated over the lifespan of the cells [6].

## Results and analysis

### Predicted volume effects of PIEZO1 activation during and after capillary transits

The four panels of figure 2 show results abstracted from a large number of simulations, to illustrate the effects of the different permeabilities and permeability combinations used to represent the open state of the PIEZO1 channels. Figure 2A shows the effects of increasing PzCa at a constant elevated PzA. At PzCa = 0, RBC volume remains at baseline level. PzCa values > 0 elicit a biphasic response a sharp initial swelling followed by slow shrinkage towards below baseline levels. Increasing PzCa leads to higher initial peaks followed by faster and deeper dehydrations. In figure 2B the Ref pattern with PzCa = 70h^−1^ and PzA = 50h^−1^ was followed for 60 minutes to expose the extremely slow reversibility from post-transit dehydration back towards baseline volume level. Without a simultaneous increase in anion permeability (PzA = 0; Fig 2C, red), the swelling response to PzCa is markedly reduced thus exposing a powerful rate-limiting effect of the anion permeability on the peak response. Attribution of sodium and potassium permeabilities to PIEZO1, PzNa and PzK in Fig 2D (red), at the upper limit of measured values in on-cell patch-clamp experiments [Dyrda], caused only a minor parallel displacement upward on the Ref pattern. These results validate the exclusion of these permeabilities from the minimalist representation of the PIEZO1 effects in the chosen reference pattern, focussing all the attention on the two critical permeability components of the Ref response, PzCa and PzA.

The most important conclusion from the predictions in Fig 2 is that all the PIEZO1-mediated effects on RBC volume during capillary transits follow from increased PzCa. PzA, PzNa and PzK modulate the response (Figs 2C and 2D), but without PzCa there is no response. PzA controls the magnitude of the peak response and is therefore the second most important component of the PIEZO1-mediated effects in the capillary context. This is particularly relevant to uncertainties concerning whether or not PIEZO1 is the unique mediator of both cation and anion permeabilities recorded in different experimental setups [27, 46−48].

Obviously, model simulations cannot resolve this issue, but what they do show is that the coordinated activation of both permeabilities is critical for the optimized magnitude of the peak response, a suggestive argument for unique PIEZO1 mediation.

Three unexpected insights emerge from the predictions in Fig 2: Firstly, the infinitesimal magnitude of the PIEZO1-induced relative cell volume changes (< 6*10^−5^); Secondly, their effective irreversibility within likely inter-transit times (Fig 2B), and Thirdly, a kinetics dominated by sharp swelling during the narrow transits and by shrinkage after egress (Fig 2A), a seemingly paradoxical sequence for smooth flow through capillaries if the volume displacements are perceived as fulfilling a mechanical role, an impossible assignment for such miniature displacements.

### Mechanisms behind the predicted volume effects

We focus here on the reference condition (black curves) unless stated otherwise. PIEZO1 activation triggers CaCl_2_ influx driven by the large inward electrochemical calcium gradient. CaCl_2_ influx elevates cell osmolarity (COs) causing a sharp fall in medium osmolarity (MOs) (Fig 3A) thus generating a gradient for water influx into the cells (Fig 3B). After PIEZO1 closure and capillary exit, medium composition is instantly restored (vertical segments in Fig 3), simulating the return of the RBC to the systemic circulation. This causes a sharp reduction in the osmotic gradient, but not yet its disappearance. Although the water permeability of the RBC membrane is very high, the magnitude of the CaCl_2_-altered osmotic gradient is miniscule, hence water influx persists for a few more seconds (Fig 3C) shaping the peak of the volume response to PIEZO1 activation (Fig 2A) beyond the open state duration of PIEZO1 and before all the downstream effects of CaCl_2_ influx take over the slow dehydration phase of the biphasic response.

**Figure 3.**
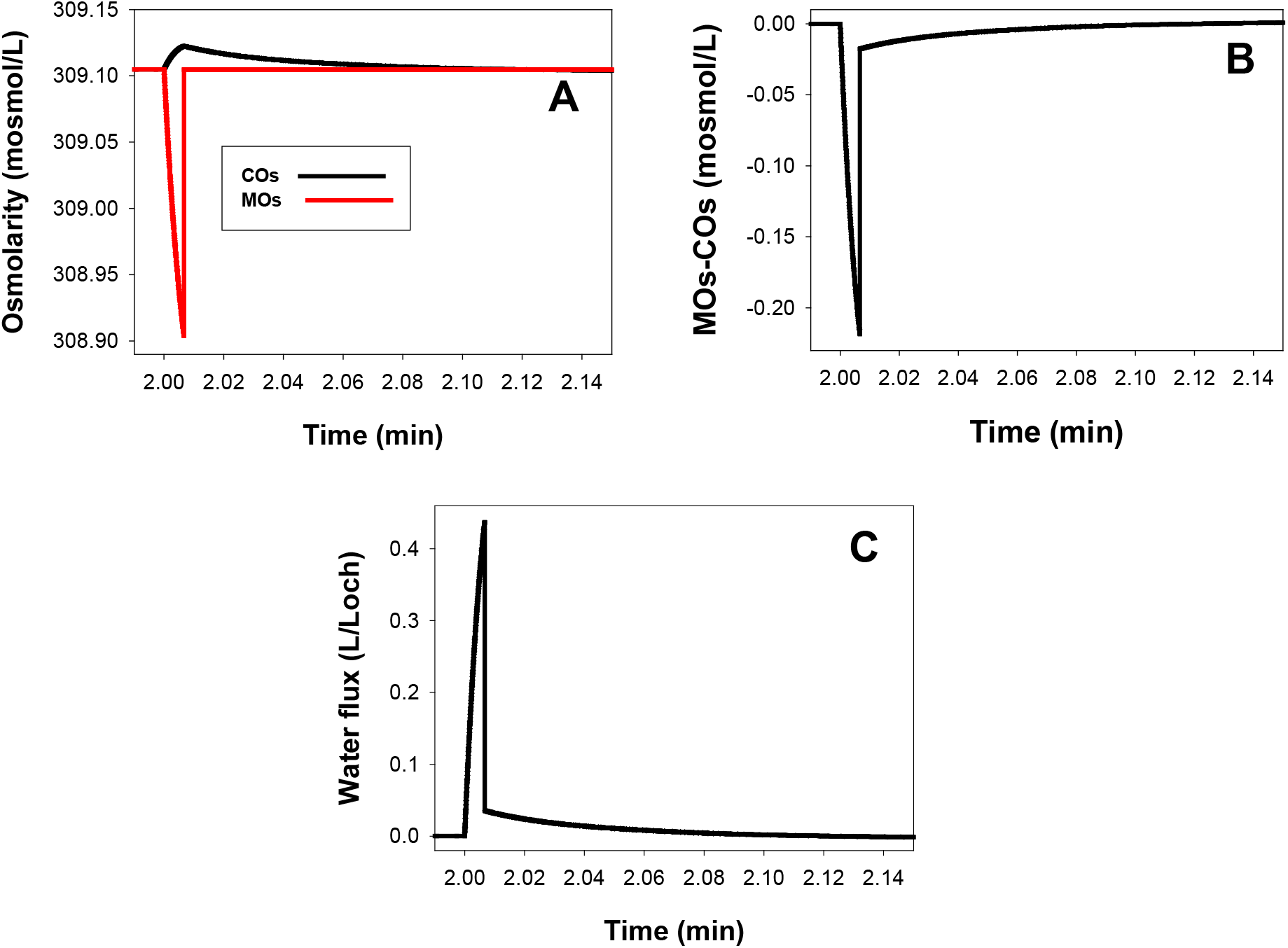
Predicted changes in osmolarity (A), osmotic gradient (B) and water flux (C) across the RBC membrane during a capillary transit modelled following the reference protocol. PIEZO1 activation triggers CaCl_2_ influx increasing cell osmolarity (COs) during the 0.4s duration of the open state and capillary transit. With a cell/medium volume ratio of 0.9 during this period, the salt shift from medium to cell caused a sharp fall in medium osmolarity (MOs) generating a gradient (B) for water influx (C) into the cells. After PIEZO1 closure and capillary exit, medium composition is instantly restored (vertical segments), simulating the return of the RBC to the systemic circulation, causing a sharp reduction in the osmotic gradient (B). Water influx persists for a few more seconds (C) shaping the peak response to PIEZO1 activation (Fig 2A), before all the downstream effects of CaCl_2_ influx take over the slow dehydration phase of the biphasic response. Note the miniscule overall mag exp

The four main transport systems involved in the predicted volume response during capillary transits are PIEZO1, Gardos channels, PMCA, and the Jacobs-Stewart system (JS). The fluid flows contributed by each are indicated in the diagram of Fig 4 by the wide arrows W1, W2, W3 and W4, respectively, and further analysed in Fig 5. Figure 5A reports the effect of Gardos channel inhibition. It shows an elevated peak response followed by shrinkage towards the baseline level, never below. Any dehydration beyond baseline is therefore entirely the result of Gardos channel activity via W2 (Fig 4). Without Gardos channel activity, dehydration mediated solely by the PMCA (W3) and the JS (W4) mechanisms only accomplish restoration of the calcium, chloride and water initially gained via W1. The elevated peak response with Gardos channel inhibited (Fig 5A, red relative to Ref) indicates that early Gardos channel activity (W2) can also antagonize slightly the peak-swelling led by W1.

**Figure 4.**
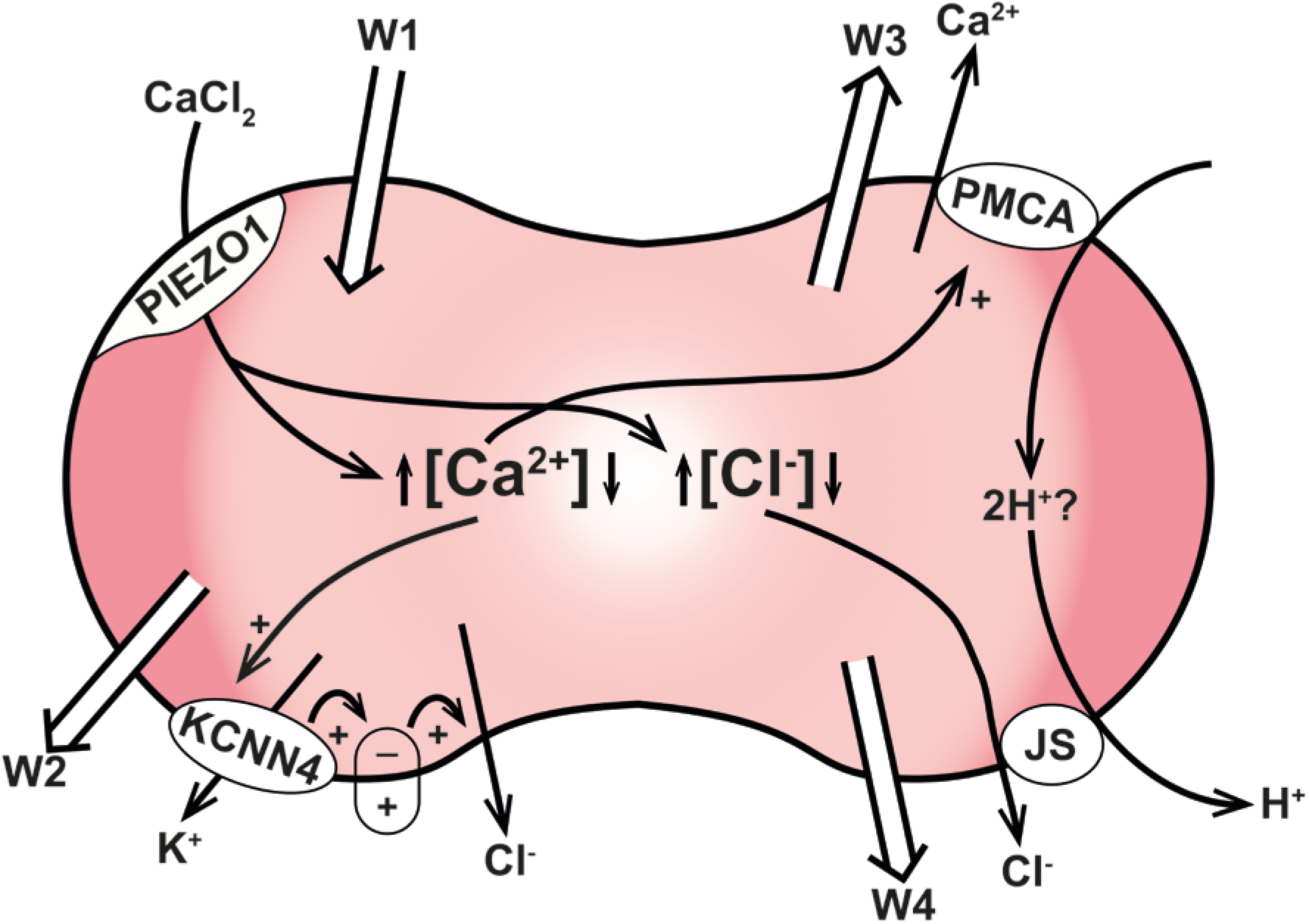
Red blood cell volume control during capillary transits. The four main transporters involved in transit-intertransit fluid dynamics are PIEZO1 (W1), KCNN4 (W2, Gardos channels), the PMCA (W3) and the Jacobs-Stewart mechanism (W4, JS). The white wide arrows indicate the direction of fluid flows during capillary passages. PIEZO1 activation triggers CaCl_2_ and fluid gain (W1) leading to a tiny but sharp increase in cell volume and [Cl^−^]_i_, and a much larger relative increase in [Ca^2+^]_i_. Elevated [Ca^2+^]_i_ activates Gardos channels (W2) and the PMCA (W3). Elevated [Cl^−^] stimulates HCl and fluid loss through the JS mechanism (W4). Gardos channel activation induces KCl and fluid loss (W2, Fig 5A). The PMCA, represented here as an electroneutral Ca^2+^:2H+ exchanger [57–59], extrudes the Ca^2+^ gained via PIEZO1 concomitantly elevating [H+]_i_ which stimulates further HCl extrusion via JS (W4). The PMCA restores [Ca^2+^]_i_ to pre-transit levels within seconds of PIEZO1 closure (Fig 5C), setting the duration of the period during which dehydration is dominated by Gardos channel activity (W2, Fig 5D). The rate of JS-mediated dehydration (W4) depends on the proton and chloride concentration gradients and on the value attributed in the model to the JS turnover rate (Fig 4E), with a default Ref value based on early measurements [Carol 87]. Fluid balance at each instant of time is set by the relation between W1 and the (W1+W2+W3) sum.

**Figure 5.**
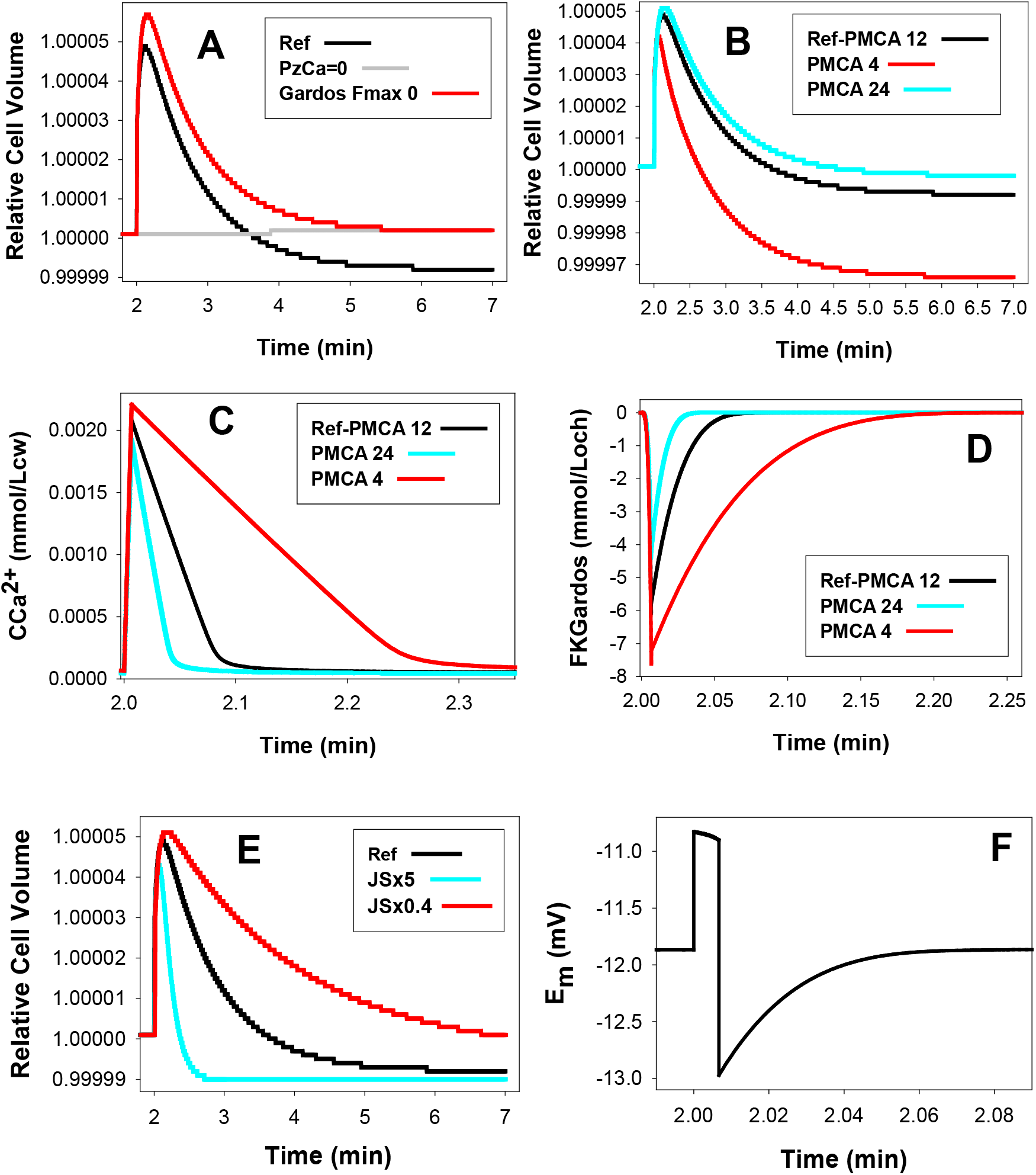
Predicted effects of variations in the activities of Gardos channels, calcium pump and JS-mediated ion transport during capillary transits. All effects reported relative to Ref (black curves). Note differences in x-axis time scales. **A.** Volume effects of setting FKmax though Gardos channels to 0 in Reference State. **B, C and D:** Effects of setting the PMCA FCamax values between 4 and 24 mmol/Loch in the Reference State on RCV (**B**), on [Ca^2+^]_i_ (**C,** CCa^2+^), and on potassium efflux through Gardos channels (**D**, FKGardos). **E.** Volume effects of reducing the default JS rate by 60% (JSx0.4) or of simulating it fivefold (JSx5). **F.** Ref membrane potential changes during the ~5 seconds following PIEZO1 activation. Note that depolarization is sustained only during the 0.4s period of CaCl_2_ influx through PIEZO1. After PIEZO1 closure the membrane potential immediately reverts to Gardos channel induced hyperpolarization, with a recovery kinetics set by Ca^2+^ extrusion and Gardos channel deactivation.

### Effects of PMCA strength on the pattern of volume change

Figures 5B to 5D report the predicted effects of different PMCA strengths on the PIEZO1 volume response (Fig 5B), on [Ca^2+^]_i_ levels (Fig 5C), and on calcium induced K^+^ flux through Gardos channels (Fig 5D). The volume curves (Fig 5B) show that the weakest the pump the lowest the peak height and the fastest and more profound the subsequent dehydration, a pattern explained by the time-course of changes in [Ca^2+^]_i_ (Fig 5C) and of Gardos-mediated K^+^ fluxes (Fig 5D). Fig 5C shows that a weak pump allows more calcium influx initially and takes longer than stronger pumps to extrude the calcium gained via PIEZO1. The three effects that result from enhanced Gardos channel activity in cells with weak calcium pumps are a larger initial Gardos-mediated K^+^ efflux (Fig 5D), a reduced volume peak (Fig 5B), and a faster and deeper post-transit dehydration (Fig 5B).

According to Figs 5C and 5D, PMCA and Gardos channel activities are back to baseline within less than 20 seconds after PIEZO1 closure, even for the weakest pumps. This means that all dehydration mediated by the W2 and W3 components is completed during this period. Yet, the volume effects become apparent only minutes later, when the initial chloride gained during W1 has been fully restored to the medium via the JS cycle (W4). The comparative volume effects of changes in JS rate are shown in Fig 5E. It can be seen that the time it takes to expose Gardos-induced dehydration is determined by the displaced chloride concentration gradient and the JS transport rate (W4). These results show that the extent of dehydration elicited by Gardos channel induced KCl loss has to await the full extrusion of the chloride load to be revealed (Figs 5B and 5E), protons providing the Cl^−^-co-ion through JS. The JS transport rate was found to vary among RBCs from adult healthy donors with a normal distribution and a coefficient of variation of 13-15% [49], variation to be noted for the likely spread of volume restoration rates around Ref (Fig 5E).

### Influence of anion exchange (JS) and anion permeabilities (PzA and PA) on the kinetics of the volume response

Although JS dominates late volume kinetics (Fig 5E), it also has a minor effect on peak height, suggesting that during the open state and the first few seconds after W1, when W2 is the main hidden player, minor contributions from W3 and W4 are also able to influence the peak volume kinetics. It may seem puzzling that a JS rate with default values set at measured levels [50], capable of flux rates orders of magnitude larger than those of the PMCA and Gardos channels should mediate the slowest kinetic component of the PIEZO1 volume response. The answer rests with the negligible magnitude and opposite displacements in the Cl^−^ and H^+^ driving gradients, which interested readers may wish to explore further using the detailed information in the csv data files.

The rate at which Gardos-mediated KCl loss and dehydration (W2) proceeds during the few seconds after PIEZO1 closure is much reduced relative to the that during the brief open state period. This is because Ref PzA (50h^−1^) was set at about fifty fold higher than PA (1.2h^−1^), the ground elecrodiffusional anion permeability of the RBC membrane [27, 38, 50–54]. Hence, after closure, Gardos-led dehydration becomes significantly more rate-limited by the anion permeability than during the brief open state, yet most of W2-led dehydration takes place post-closure because of the extended period of elevated [Ca^2+^]_i_ (Fig 5C).

### Membrane potential changes during capillary transits

Figure 5F shows the membrane potential changes predicted for the first five seconds of the reference protocol following PIEZO1 activation. The initial depolarization during the Ca^2+^ influx-dominated open state changes abruptly to Gardos-dominated hyperpolarization after closure, with a relatively slow return to baseline membrane potential following the calcium desaturation kinetics of the Gardos channels. The full magnitude of the predicted membrane potential displacement was about 2 mV, a miniaturized version of that fitting the results of the on-cell voltage clamp experiments with a much delayed PIEZO1 inactivation kinetics [27].

## Discussion

The results presented here offer the first exploration of the behaviour predicted for human RBCs during single capillary transits applying a realistic representation of the process in compliance with available knowledge. Model outputs provided a detailed high-resolution account of the changes in RBC homeostasis variables during and following capillary transits, changes triggered by deformation-induced activation of PIEZO1 channels on ingress of RBCs into capillaries. Analysis of the results and interpretation of the mechanisms involved was followed on large sets of simulations, summarily reported and analysed on figures 2 to 5.

The main features of the predicted volume response of RBCs to a capillary transit initiated by PIEZO1 channel activation were i) a biphasic sequence of rapid initial swelling followed by slow shrinkage towards below-baseline volume levels (Fig 2A), ii) extremely slow post-transit reversibility (Fig 2B), and iii) the infinitesimal magnitude of the predicted volume displacements, incompatible with functional role assignments on capillary blood flow (Figs 2 and 5). The four main mechanisms involved in osmolite and fluid transfers during and following capillary transits are depicted in the diagram of Fig 4. PIEZO1 is responsible for initial swelling driven by CaCl_2_ gain. The PMCA restores calcium fully (Fig 5C); Gardos channel-mediated K^+^ efflux causes a miniscule potassium depletion (Fig 5D), and the Jacobs-Stewart mechanism (Fig 5E) restores chloride and pH towards new levels when dehydration settles below baseline volume levels, strictly the result of early Gardos channel activity (Fig 5A). The time-lag to the final dehydrated quasi-steady-state is largely determined by the chloride efflux rate through the Jacobs-Stewart mechanism (Figs 2B and 5E).

The main conclusions concerning the quantal hypothesis are that volume changes during single capillary passages have the potential to progressively accumulate, densify, swell or balance RBC volumes in the circulation depending on whether the duration of the inter-transit periods allows net dehydration, net swelling or balanced volume change to prevail (Figs 5B and 5E) during the changing homeostatic conditions of aging RBCs, alternatives to be investigated and tested in the following paper [6].

Danielczok et al., [35] suggested a Ca^2+^-mediated adaptation mechanism in which RBCs were supposed to experience relatively large and rapidly reversible volume reductions during capillary transits in aid of smooth capillary flow. This hypothesis was based on a perceived need for RBCs to be superdeformable on entering capillaries, for which a transient and sizeable initial volume decrease was considered necessary. GsMTx, a known inhibitor of PIEZO1 channels [55, 56] was found to cause RBC shape changes, microfluidic block and reduced RBC filterability. Interpreting these effects as the result of PIEZO1 inhibition preventing the initial volume reduction necessary for transit, the GsMTx effects were considered supportive of the adaptation hypothesis. However, altered RBC filterability is not the normal condition of RBCs with normally silent PIEZO1 channels, suggesting that GxMTx or the GsMTx-PIEZO1 interaction, besides channel block, induced additional abnormal side-effects in the conditions of their experiments, of doubtful relevance as tests of the adaptation hypothesis. In addition, as analysed in relation to figures 2 and 5 here, the first volume deflection induced by PIEZO1 channel activation is swelling, not shrinkage. Above all, the known constitutive properties of human RBCs do not allow volume changes of the magnitude, speed and reversibility time-course required for the kind of flow dynamics function required by the adaptation hypothesis.

## Acknowledgements

The background research work on which this investigation was based was supported over four decades by funding agencies in the UK and in the USA (VLL PI or co-PI). **In the UK:** Biotechnology and Biological Sciences Research Council (BB/E008542/1; BB/F001630/1, BB/F001673/1, and BB/H024867/1), Engineering and Physical Sciences Research Council (EP/E059384); The Wellcome Trust (064124; 061269; 059725; 030699; 033876; 15543; 17358; 13056), and The Medical Research Council (G8211073CA). **In the US:** NIH 2-RO1 HL28018-19; 2-RO1 HL21016-11. The authors are grateful to Serge L. Y. Thomas and Daniel J. Lew for helpful comments and suggestions on the material contained in the three manuscripts of this series.

